# Social migratory connectivity: do birds that socialize in winter breed together?

**DOI:** 10.1101/2021.12.03.471164

**Authors:** Theadora A. Block, Bruce E. Lyon, Zachary Mikalonis, Alexis S. Chaine, Daizaburo Shizuka

**Affiliations:** Department of Ecology and Evolutionary Biology, University of California, Santa Cruz, California, United States of America; Station d’Ecologie Théorique et Expérimentale du CNRS (UMR5321), Evolutionary Ecology Group, Moulis, France; Institute for Advanced Studies in Toulouse, Toulouse School of Economics, Toulouse, France; School of Biological Sciences, University of Nebraska-Lincoln, Nebraska, United States of America

## Abstract

Researching the complete life cycles of migratory animals is essential for understanding conservation and population dynamics. Many studies focus on the breeding season, but surviving winter is equally important. Living in groups during winter can play a vital role as social connections within groups can provide many benefits such as protection from predators and increased access to resources. However, it is often unknown how social connections change across seasons in migratory animals. We focus on social connections in a migratory bird and ask whether social connections in winter continue during breeding. Golden-crowned sparrows have distinct, stable winter communities which include both site and group fidelity across years: birds almost always rejoin the same social community each year. If these birds have social connectivity across migration, we would expect individuals that associate in winter would also associate together on their breeding grounds. Our small-scale GPS tagging study combined with intensive social behavior data revealed that sparrows in the same tightly-knit winter community migrated to highly disparate locations during summer, showing that social connections in winter do not continue in summer. This suggests that golden-crowned sparrows have entirely separate social structures across seasons and that long-term social memories allow them to reform stable groups each winter.

## Introduction

The study of migratory connectivity seeks to understand how breeding and nonbreeding populations of migratory animals are functionally linked, generally focusing on migratory patterns of populations across broad regions [1, 2]. Discovering migration patterns can be vital to understanding population dynamics and conservation [1, 3, 4–7]. For example, both golden-winged warblers (*Vermivora chrysoptera*) and Swainson’s thrushes (*Catharus ustulatus)* have strong migratory connectivity, where populations from one breeding ground go to the same nonbreeding area – hence, their overall population declines are linked to degradation of specific locations [8, 9]. However, migratory connectivity could operate on a finer scale if individual-level social connections between migratory animals persist from nonbreeding to breeding seasons. We define social migratory connectivity as social connections between migratory animals present in one season and maintained in the next. Social migratory connectivity focuses on migration at the level of individual connections and requires geographic precision and detailed information about social interactions. If present, social migratory connectivity could fundamentally affect how social relationships are formed and maintained in migratory animals. These relationships cannot be fully understood by observing them in only one part of the year.

For example, do social relationships in one season carry over to affect social relationships in another season? Such carry-over effects are seen in year-round resident birds such as great tits (*Parsus major*), where connections in winter predict associations in the breeding season or in blue tits (*Cyanistes caeruleus*), where social connections in winter predicted nesting proximity and even increased extra-pair paternity [10, 11]. In theory, such social carry-over effects could exist in migratory birds as well, but this idea is rarely tested. In one remarkable case, migratory European bee-eaters (*Merops apiaster*) maintain cohesive social relationships across seasons by migrating together [12]. Additionally, pine siskins (*Carduelis spinus*) can form stable groups throughout the year, traveling distances over 1000 km together, and in some cases, groups stayed together for almost four years [13]. Some other social relationships within one season have spanned across years in migratory birds (e.g., ‘dear-enemy’ effects among territorial neighbors in breeding season: [14]; flock mate relationships in winter: [15]), but we currently do not know if these represent carry-over effects of social relationships that were established in other seasons.

Long-term research on golden-crowned sparrows gives us a sufficiently nuanced understanding of their winter ecology and sociality to test whether social connectivity is maintained across seasons. Golden-crowned sparrows (*Zonotrichia atricapilla*) are small migratory birds that live in complex societies during winter. The sparrows winter in relatively stable communities with high site fidelity across years, and associations are based on social preference more than overlapping space use [15]. Birds that return to the winter site almost always return to the same social community and tend to increase the strength of their associations with other individuals present [15]. While there are related individuals in the population at the wintering grounds, genetic kinship does not predict patterns of association [16]. This raises questions about how sociality might change across seasons and whether migration and breeding location could connect the birds’ social patterns across seasons. If golden-crowned sparrows have linked breeding and nonbreeding grounds, we might also expect that birds with close winter social relationships might have more social interactions during the breeding season. Golden-crowned sparrows breed in Alaska and western Canada in shrub habitat that is near timberline or in coastal areas [17, 18], where they form socially monogamous pairs (the extent of extrapair mating has not been studied). Social interactions during the breeding season consist mainly of social mates and agonistic interactions between adjacent territory holders, often through vocal communication (e.g., [19]). Evidence from other species suggests that young animals make social connections with neighbors early in life; for example, captive barnacle geese (*Branta leucopsis*) formed connections early in life and carried these same preferences into adulthood [20] and across seasons [21]. Currently, we know that golden-crowned sparrows have broad-scale migratory connectivity, as birds wintering in coastal California tended to go to coastal areas in Alaska while more inland birds went to more inland areas in the north of Alaska to breed [22, 23].

Here, we ask if golden-crowned sparrows have social migratory connectivity, or, if they maintain close social connections from the nonbreeding season during the breeding season. Two main patterns are possible: 1) golden-crowned sparrows from the same winter social communities could breed in close proximity, or 2) winter associations may be entirely unconnected to breeding locations and associations. The first pattern would reveal that the sparrows maintain social connections year-round and strongly suggest that associations during the breeding season and first migration are critical for establishing and maintaining the close social associations we observe in these birds during winter. The second pattern would be equally interesting as it would reveal that these birds have a strong capacity to remember individuals and have long social memories to reform stable communities each winter.

We used archival Global Positioning System (GPS) tags to study golden-crowned sparrow social migratory connectivity. GPS tags light enough to use on small birds are a relatively new technology, and researchers have only been able to collect this sort of precise location data across seasons for a few species of songbirds (see: [9, 24–26]). GPS tags open opportunities to discover exact migration paths and breeding locations. This technology gives us the geographic resolution to answer questions about how social connections change across seasons. While we report a small sample size, we have detailed data on individual social interactions prior to migration and follow birds from the same winter social group. In addition to social migratory connectivity, the suggestive patterns we observed motivate further questions about the relationships we might expect between migration departure dates, duration of migration, and time spent on summer breeding grounds.

## Methods

Golden-crowned sparrows arrive at our field site at the University of California Santa Cruz (UCSC) Arboretum (36.9841, -122.0599) around the end of September. They form winter flocks and remain at the study site until April/May when they leave for their breeding grounds in northern Canada and Alaska. Our study focused on the migratory patterns of individual golden-crowned sparrows after observing the birds for at least one season on their wintering ground. As part of an ongoing larger study, we gathered morphological and behavioral data from golden-crowned sparrows during their nonbreeding season at the UCSC Arboretum. We caught, measured, and banded the sparrows and collected a blood sample from the ulnar vein to sex individuals [27, 28]. During banding, each bird received a U.S. Geological Survey uniquely numbered metal band and color bands with a unique color combination to be identifiable in the field.

### GPS methods

We programmed GPS locators (1 g PinPoint 10 Swift GPS tags made by Lotek Wireless) to take up to 100 locations. The length of time a tag took to read satellite signals at each point affected how long the battery life lasted, and tags were expected to record around 75-80 points. These locations are precise GPS coordinates, accurate to within 10 m (Fig 3), which gave us the precision to see if birds not only bred nearby each other, but if they had neighboring territories (golden-crowned sparrow breeding territories are approximately 0.86 ha, D.S. unpublished data). We programmed the tags to take points along the birds’ migration paths and points on the breeding ground to establish the exact location of breeding territories. The tags took points every two to five days, depending on our estimations of when the birds might be on migration versus established on the breeding grounds.

In the spring of 2017, we attached 30 archival PinPoint 10 Swift GPS tags to previously banded golden-crowned sparrows that had been at the Arboretum for at least three or more months prior or had returned from previous years. In the spring of 2018, we attached 40 GPS tags to a different sample of sparrows that met the same criteria. We used a leg-loop attachment method [29] with 0.7 mm stretchy jewelry cord. After attaching the GPS tags, we monitored sparrows in the field, ensuring that birds maintained their normal behavior and could move, fly, and feed unhindered.

Archival GPS tags must be recovered to retrieve the data, so we depended on the tagged birds returning to the UCSC Arboretum after the breeding season. Despite deploying 70 GPS tags, we only obtained adequate data from four birds over two field seasons due to several unanticipated obstacles. From 30 GPS tags in 2017, we recovered five the next field season. The low return rate of tags was due to problems with the GPS harness attachment, as at least five birds lost their harnesses before migration, and of seven previously tagged birds that returned, two had lost their tags after leaving the winter site. Of the five that returned with tags, two tags malfunctioned and gathered incomplete data (8 and 15 locations respectively, all before the birds left California), leaving us with data for three birds. From 40 tags attached to sparrows in 2018, we recovered one tag. In 2018, a substantial portion of our study site was destroyed to make a parking lot (0.54 ha) within a month of tag-deployment, in the precise location where we had focused our tagging effort. Additionally, during fall migration in 2018, multiple large fires along the California coast may have interfered with birds returning. These two factors in 2018 likely resulted in the very low return rates of tagged birds in the fall. The four tags with full data recorded 75 GPS points for bird 77968, 68 points for bird 81319, 85 points for bird 81324, and 49 points for bird 19388 [30].

We used the GPS data to determine the location of the golden-crowned sparrows’ breeding territories by searching for highly localized clusters of more than five GPS points at the furthest northern location of a bird’s migration route. The breeding locations for all four birds had a high density of GPS points over a small area (< 0.4 km^2^), and we used these locations to determine the start and end date of the time each sparrow spent on its putative breeding territory. We determined the average breeding territory location by calculating a centroid location from the clustered points using the package *geosphere* [31]. To calculate the distance of spring migration, we used the great-circle-distance between two points from the Vincenty ellipsoid method [31].

We excluded two outlier GPS points during the breeding season (one for bird 77968, one for bird 81324) that were deemed erroneous due to their extreme distance from other points clumped on the breeding grounds (327 and 615 km away) and a reduced number of satellites for those GPS fixes (3 satellites, compared to 5 or more for most other points).

### Social network methods

As part of our fieldwork during the nonbreeding season, we found which birds flocked together in the UCSC Arboretum with frequent censuses throughout the winter season. This included identifying all color-banded sparrows and marking their location from a gridded map with 10 m^2^ cells. From this data, we built social networks following methods from Shizuka et al. [15]. We modified these methods to include all birds, regardless of age, where Shizuka et al. [15] excluded birds from the social network analysis if they were banded for the first time that season. Our network analysis incorporated all birds seen more than five times in the field that season, including birds initially banded that year. We used flocking data from before April 1 of each year, as later in the season community structure can deteriorate closer to migration. From the flocking data, we built undirected social networks with the R package *igraph* [32]. The network is made up of individuals (‘nodes’), which are connected by lines (‘edges’) [33, 34]. We weighted edges by how often birds are seen together with a correction for how many times each bird is seen in total, called the Simple Ratio index [35]. The Simple Ratio index ranges from zero to one: if two birds are always seen together then they are connected by an edge value of one, but if they are rarely seen together, the value is closer to zero [35]. We determined bird community membership with a simulated annealing algorithm from the package *rnetcarto* [36] and used modularity to measure how distinct the communities were from each other. Communities were determined by calculating maximum modularity (*Q*_max_), which finds the highest proportion of edges that are within a community compared to edges across communities [37]. A higher *Q*_max_ indicates that individuals socialize more within their community than with individuals in other communities. We used a social network measure for individuals called within-community node strength (*z*-scores from [38]; hereafter called within-community strength) with the package *rnetcarto* and normalized scores [36]. Within-community strength is the sum of all edge weights a bird has to other individuals within its own community. Then, we normalized these scores using a *z*-score based on the distribution of association strengths within the community, so values were comparable for individuals in different communities and years. For example, a bird that associated with numerous individuals in its community would have a high within-community strength, as would a bird that was seen frequently with a smaller number of individuals in its community.

All analysis was performed in R version 4.0.2 [39], and maps were made in ArcMap version 10.7.1. We made a live migration track map, available in the online Supplementary Material, using the R package *moveVis* [40].

All methods complied with Federal and California State regulations under permits to B. Lyon and were approved by the UCSC Institute for Animal Care and Concern (IACUC; Animal Welfare Permit Number Lyonb1808 to B. Lyon).

## Results

All four of the tagged birds had summer locations far away from each other, with the closest 698 km apart and the most distant 1837 km apart (Fig 1). The three birds in 2017, all from the same tight-knit winter social group, had very distant breeding territories, so we found no evidence that winter community affected the proximity of summer breeding territories for these individuals. Note that the scale on which birds overlap during the winter is hundreds of meters, while in the summer, all sparrows were many hundreds of kilometers apart. Bird 19388, which we tracked in 2018, was also present on the wintering grounds at the Arboretum the previous year with the three other birds (2017), and it was in a separate social group from those three (Fig 2). The data from the GPS tags showed precise locations, with the ability to even see approximate potential nesting locations from the density of points in one location (Fig 3).

**Fig 1.**
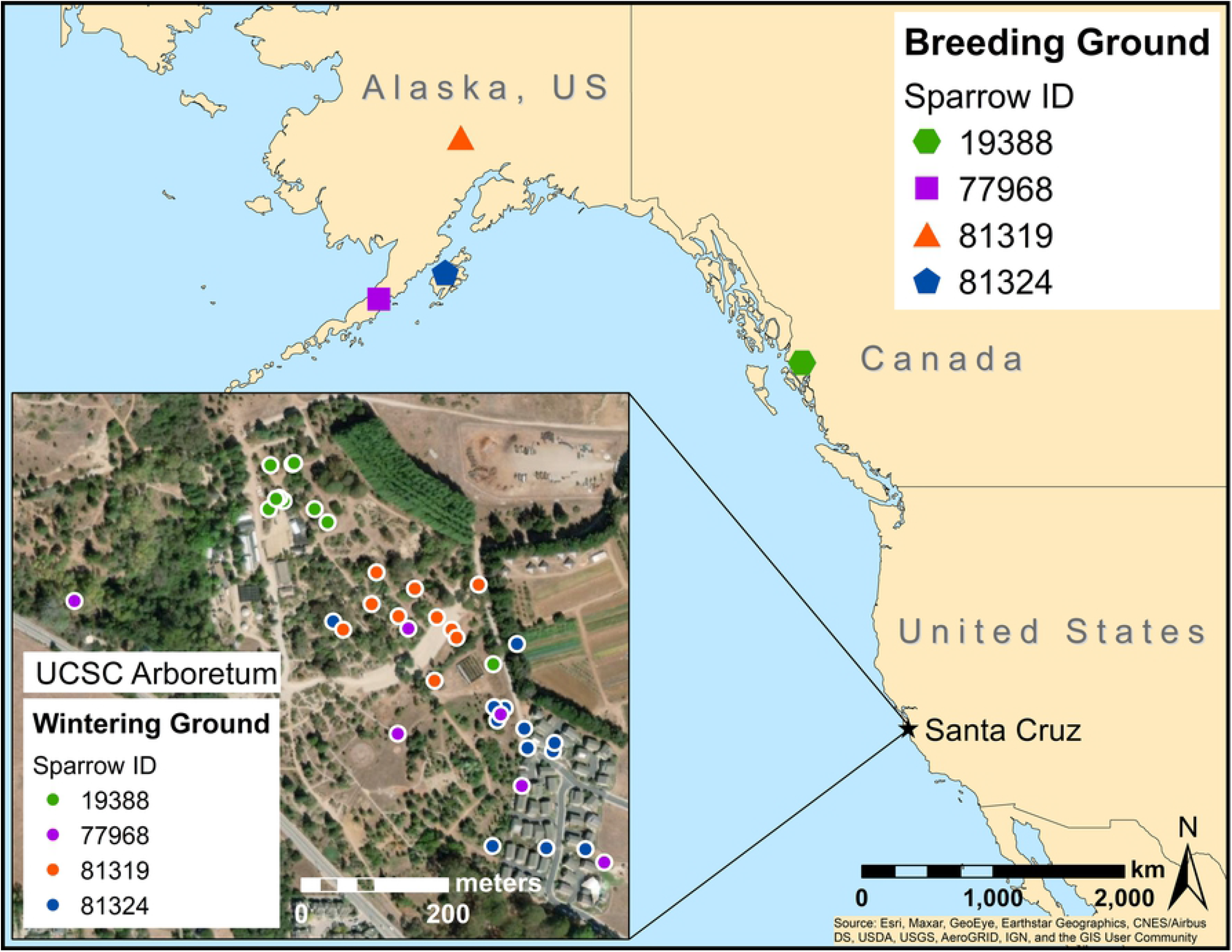
Breeding (main map) and wintering (inset map) locations for each bird. Tag data for bird 19388 was from one year after (2018) the other three birds (2017), but all birds were present in winter 2017. Symbol representation of each point is not to scale, and breeding ground areas are vastly smaller than each symbol. The points shown at the UCSC Arboretum were from the GPS tags shortly before the birds left for spring migration, not the birds’ home ranges for the winter season. To see the communities throughout winter from social network analysis, see Fig 2.

**Fig 2.**
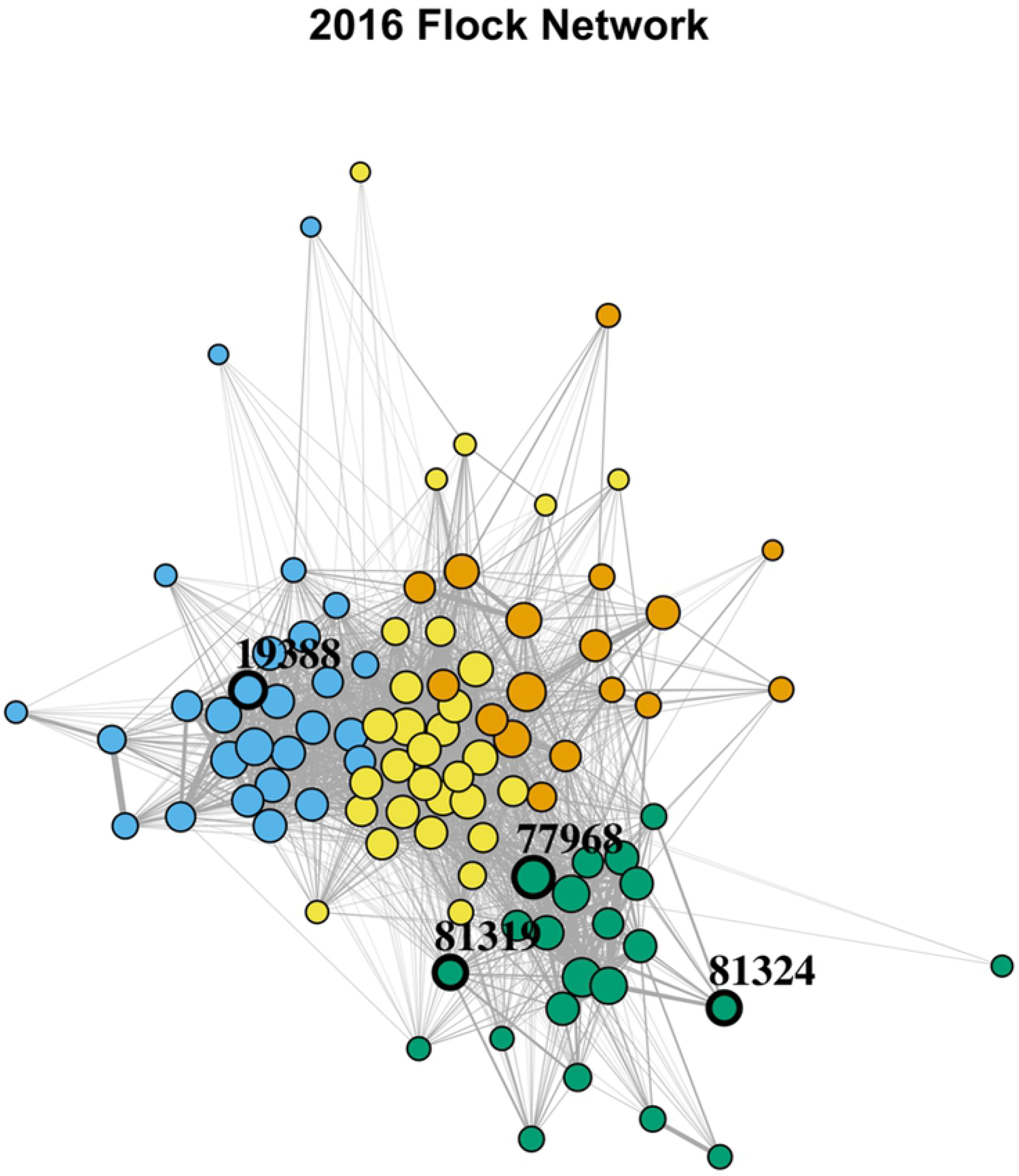
The visualized golden-crowned sparrow social network at the UCSC Arboretum for the winter of 2016–2017 (representation is not spatially explicit). The four communities are shown in different colors, and circles represent individual birds. The size of each circle is proportional to within-community strength, so larger circles show birds that have stronger associations with other birds in their communities. The thickness of the lines connecting individuals is proportional to how frequently birds associated with each other weighted by the number of times each bird was seen in total (a Simple Ratio index). The four individuals with GPS tag data are highlighted in black edging with their band ID’s. Birds 81324, 77968, and 81319 have GPS data corresponding with this year, but GPS data for bird 19388 is from the following year (2018).

**Fig 3.**
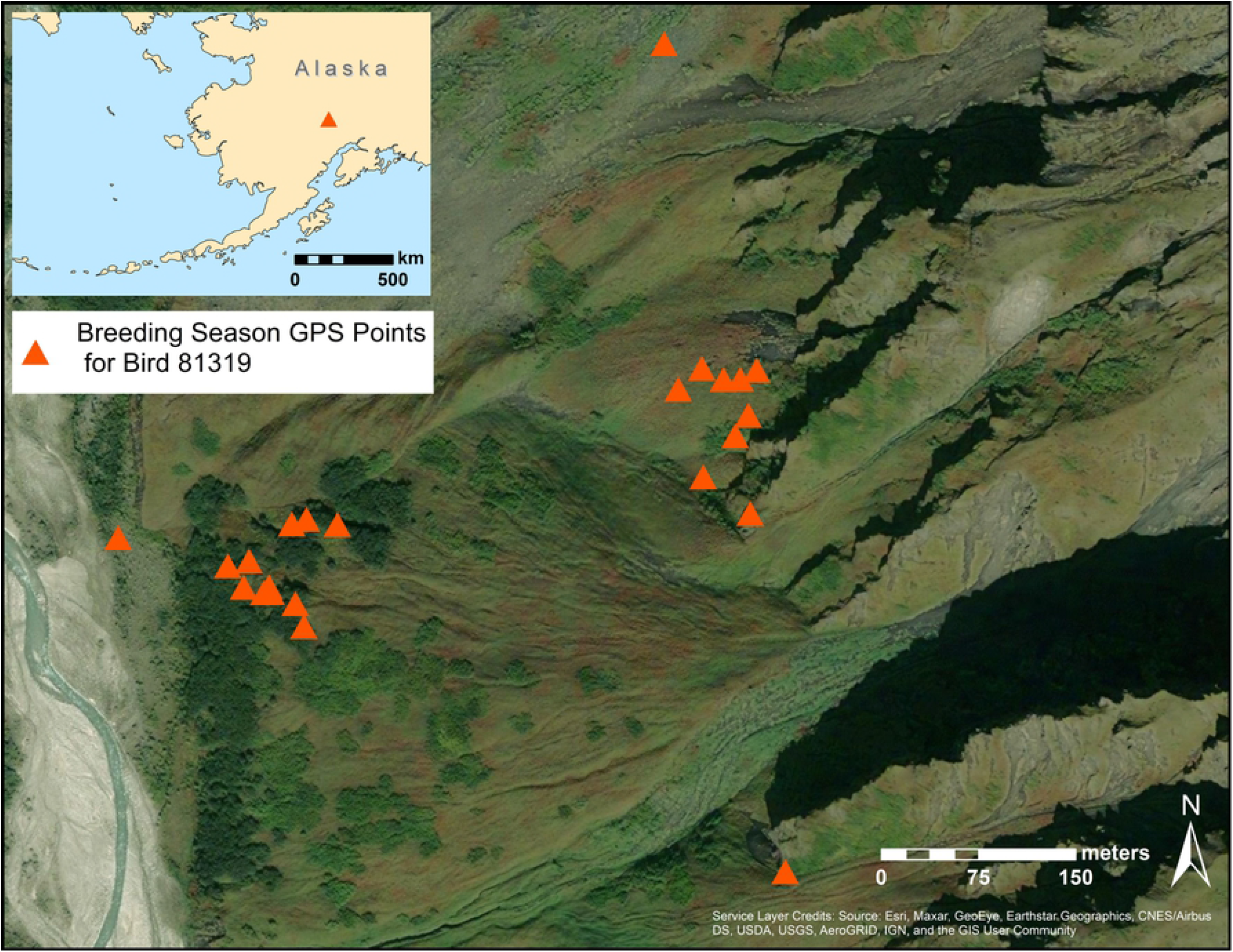
A map illustrating breeding season site occupancy and the level of resolution possible with GPS tags. Each triangle is a GPS point taken for bird 81319 on its summer breeding ground. The GPS points span June 21 to September 2 2017, the birds’ entire duration on its summer grounds. The two main clusters of points are sequential and could reflect a second nesting attempt, perhaps after nest predation, or the adult’s movement with fledglings away from the nest. We found a similar pattern with bird 19388.

The four sparrows with GPS data were representative of two different social groups. There were 96 golden-crowned sparrows in the 2016-2017 social network, with four different communities. There were 27 sparrows in the blue community, 22 in the green community, 17 in the orange community, and 30 in the yellow community (Fig 2). The four communities were distinct from each other, with a modularity (*Q*_max_ = 0.31) similar to other years of social networks in the golden-crowned sparrow system [15]. Three of the sparrows with GPS data were from the green community, representing 14% of that community.

The sparrows’ spring migrations followed the general coastline, but the birds did not use the same routes or stopover spots (Fig 5). Bird 19388 migrated further inland than the others, up to 98 km inland in parts of central California (39.10146, -122.5892) and 180 km inland in south-western Canada (50.81969, -125.8825). As this was the only bird with GPS data in 2018, it is unclear if this difference is due to a year effect or general variation among birds’ migration paths. The variation in spring migration duration appeared to originate from time spent at stopover sites rather than the number of stopovers. Most of the GPS tags stopped recording points before birds had completed fall migration, but we have the complete fall migration record for bird 81324. Interestingly, this bird spent 36 days on fall migration, identical to its 36-day spring migration (Table 1).

**Table 1.**
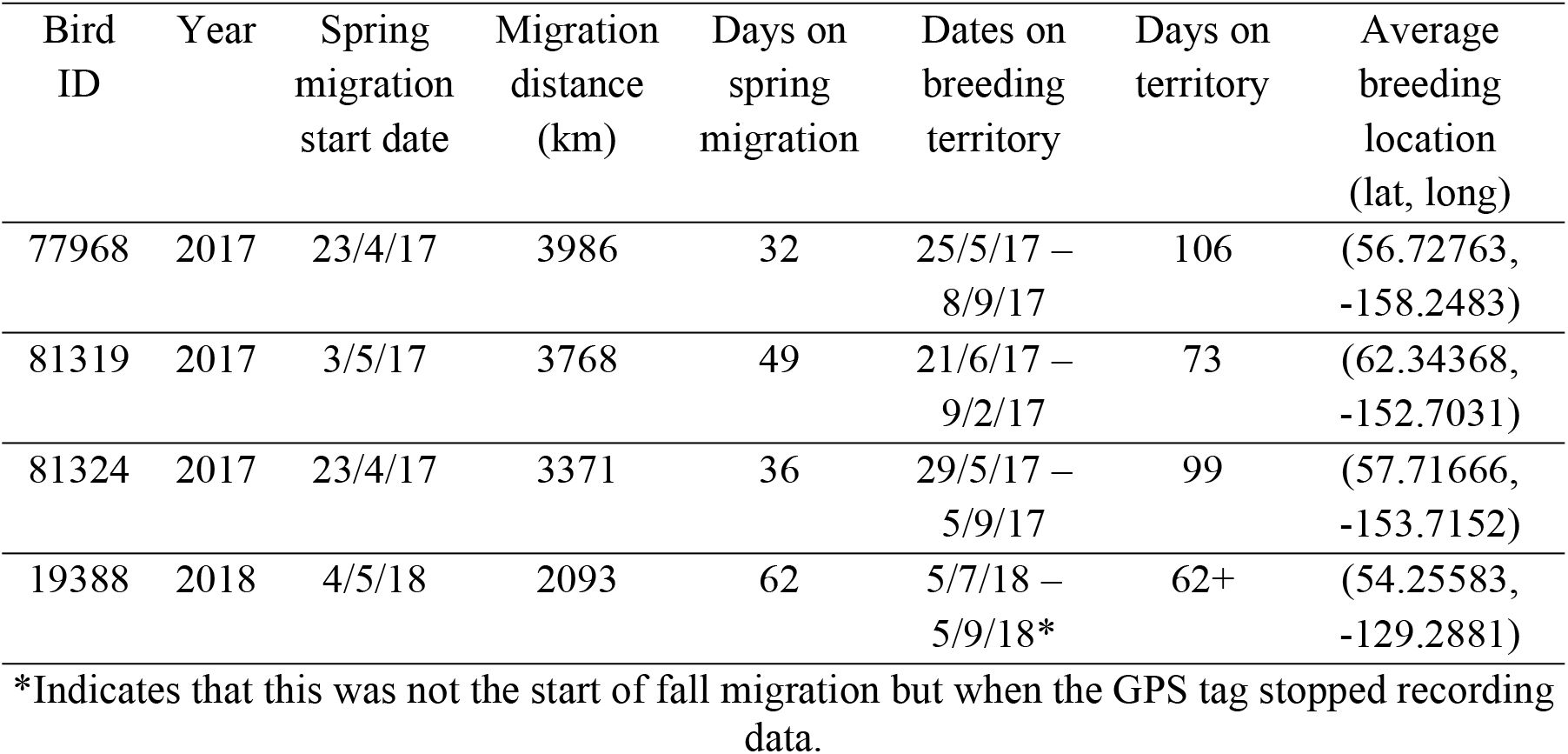
Migratory and breeding location parameters for four golden-crowned sparrows.

How quickly birds migrated made the difference in how long they spent on their breeding territory. Spring migration duration ranged from 32-62 days. The sparrows departed for spring migration from the Arboretum within 12 days of each other, and fall migration departure was within seven days of each other (Table 1). Due to such similar migration departure dates in both spring and fall, time on the breeding territory appeared to be constrained by the duration of spring migration (Fig 4). We might expect that a closer breeding location would allow the birds to get there more quickly. However, spring migration distance appeared unrelated to how many days it took the birds to migrate and even suggested a negative relationship (Fig 4).

**Fig 4.**
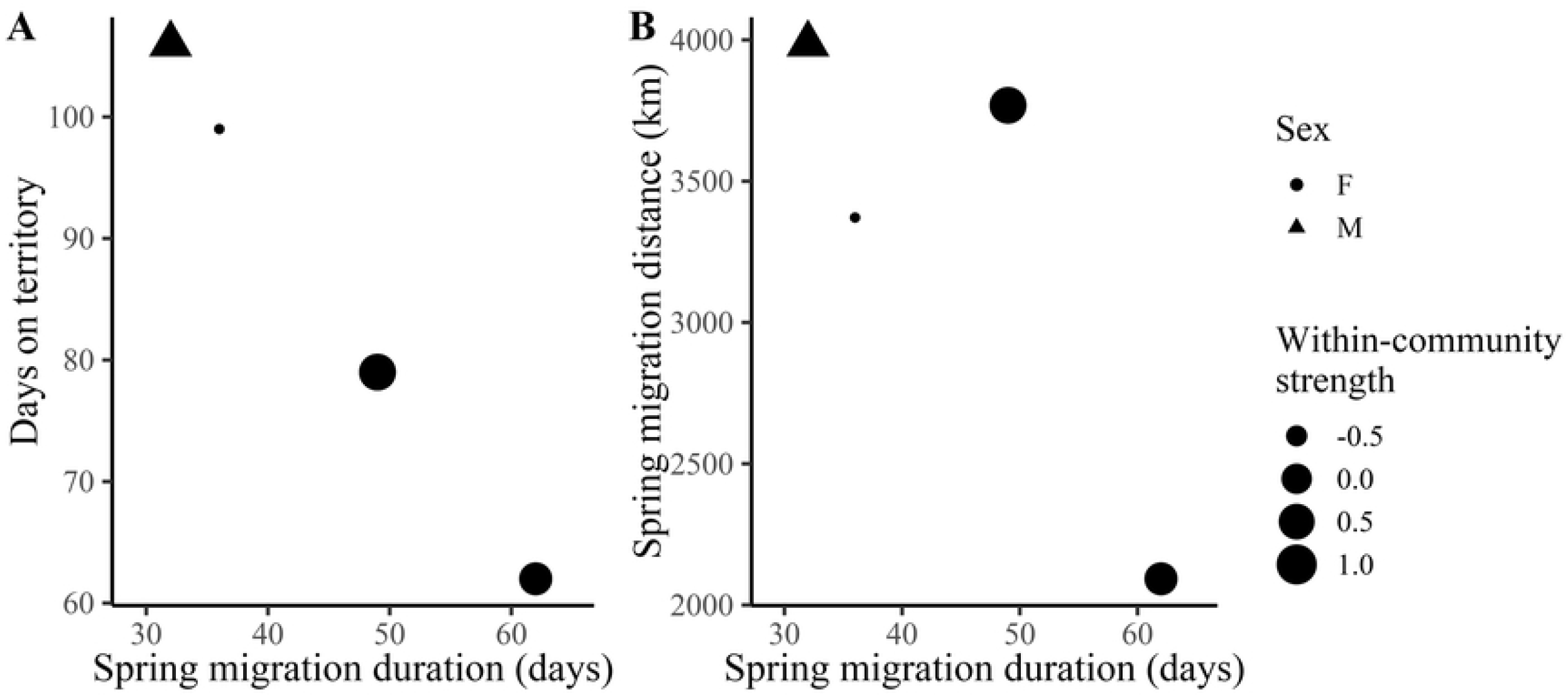
A. The number of days a bird spent on its breeding territory decreased with time spent on migration. **B**. Migratory distance appeared to have little to no relationship with how many days a bird spent on spring migration and potentially hinted at a negative relationship. Point shape indicates sex, and point size is weighted by within-community strength, or how strongly a bird associated with other individuals in its community.

**Fig 5.**
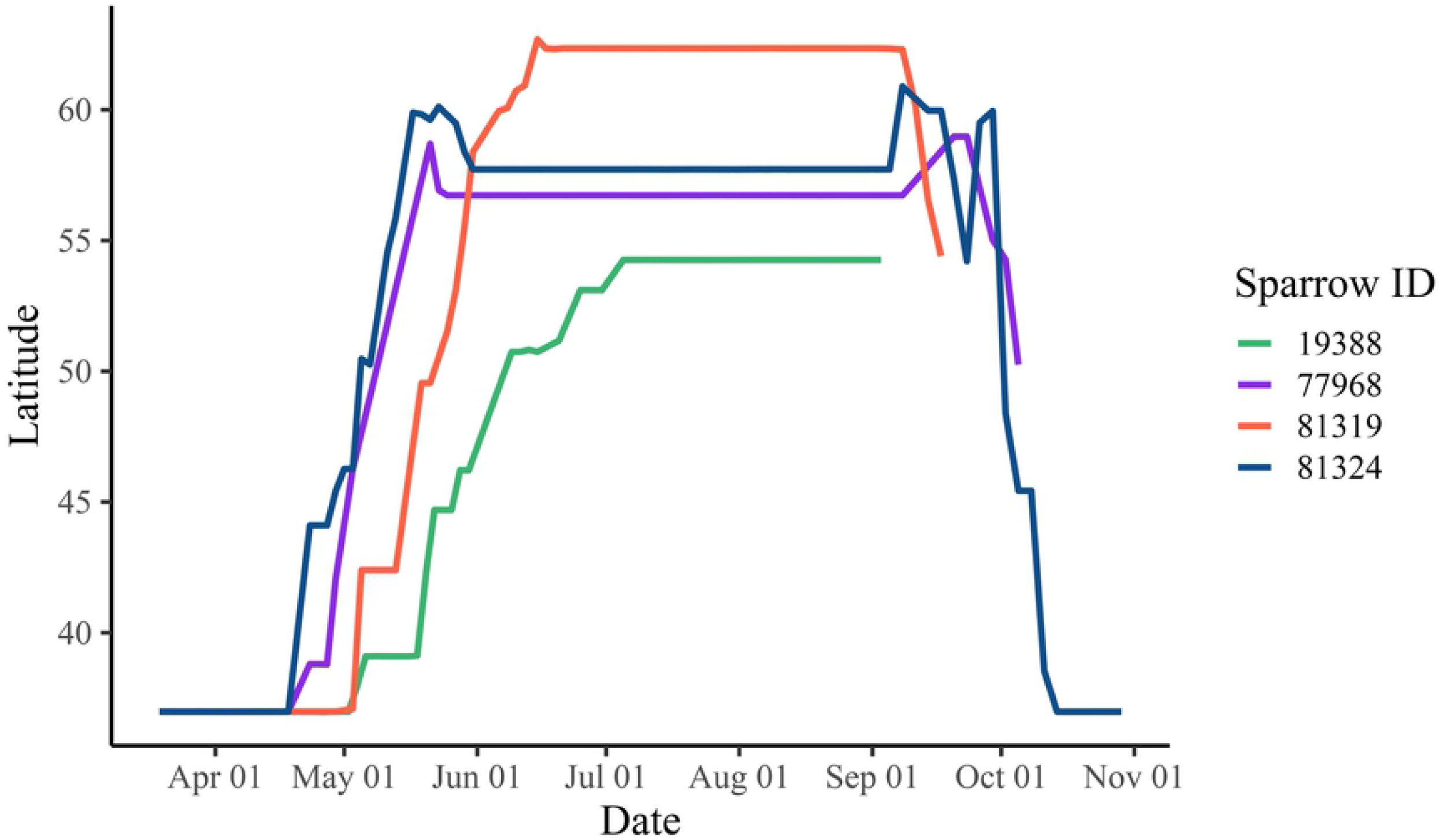
An increase in latitude from spring to fall of golden-crowned sparrows migrating north to their breeding territories illustrates the among-individual variation in the duration of migration and the location and time on the breeding grounds. The northern overshoot seen most strongly in bird 77968 (purple line) and bird 81324 (blue line) is due to migration paths following the curved coastline of Alaska (see Fig 1).

The sparrows had a wide range of within-community strength scores, but within-community strength showed no pattern when compared to migration duration or distance (Fig 4). Nonetheless, the limited sample size warrants caution on this interpretation.

## Discussion

The golden-crowned sparrows we studied all used breeding grounds that were widely separated from each other, showing entirely different social connections on winter versus summer grounds. While space use and the physical environment can play a large role in helping determine social structures [41], social affiliations and preferences are critical in driving animal interactions within groups [42]. Shizuka et al. [15] established that golden-crowned sparrows maintain social connections across years and that these social preferences are not just based on space use. This research in the same study system combined with our findings indicates that these birds have long-term social relationships and potentially long-term social memory. Other migratory birds have been shown to have social memory across years; for example, male hooded warblers (*Setophaga citrina*) recognized neighboring males’ songs eight months later during the following breeding season [14]. While we have a limited sample size for the location of the sparrows’ breeding territories, the social affiliations from the field are comprehensive. If birds in the same winter community went to similar breeding grounds, we should have seen some indication with breeding locations closer to each other, rather than hundreds of kilometers apart.

The phenology of migration can have many repercussions on the success of the breeding season and fledging young, and the variation of timing can be due to wind, weather, temperature, and photoperiod [43, 44]. The timing of the golden-crowned sparrows’ migrations revealed some interesting suggestive patterns that could motivate future investigations. Despite being part of the same winter social community, the sparrows did not appear to migrate together or use any of the same stopover sites. The golden-crowned sparrows all departed for spring migration in a relatively small window of time (12 days), so arrival on the breeding grounds is potentially determined by the speed of migration and length of time spent at stopover sites. Speed of migration is subject to many factors, whether environmental or based on individual behavior and flexibility. Similar to golden-crowned sparrows, the length of time spent at stopover sites also determined how long migration took in Northern Wheatears (*Oenanthe oenanthe*) [45]. This contrasts with wood thrushes (*Hylocichla mustelina*), where the date left on spring migration correlated with the arrival date on the breeding grounds [46]. Along with environmental factors, it could be necessary to consider individual variation in migration timing and how that affects the length of time on the breeding grounds, especially in species which have similar departure dates.

Golden-crowned sparrow winter communities are relatively stable with some turnover [15]. Communities continue over multiple years despite changing composition due to deaths and new individuals joining. How new birds integrate into these communities will be vital to understanding social group maintenance and stability over time [15, 47]. While we found no connection between winter social communities and summer locations, previous work in winter found genetic similarities in the UCSC Arboretum sparrow population [16]. Many individuals in the study population had genetic similarities (relatedness coefficient ≥ 0.25), which could be due to two main reasons: first, genetic similarity due to kinship relationships, or, second, golden-crowned sparrows may have genetically distinct breeding populations, so unrelated individuals from the same population would appear to have higher amounts of genetic overlap [16].

Consequently, either related individuals or individuals from the same breeding population may migrate from the breeding grounds to similar wintering areas. Perhaps related juveniles migrate from a shared breeding ground to the same over-winter location in their first year, then migrate to different breeding grounds in the following years while keeping fidelity to their initial winter site. This pattern has been observed in migratory pied avocets (*Recurvirostra avosetta*) and greater flamingos (*Phoenicopterus roseus*), both of which show high fidelity to their first wintering site despite differing breeding sites [48, 49]. Hence, low natal breeding philopatry does not require low wintering site fidelity.

Three of the GPS-tagged birds were in the same community (Fig 2), so if close winter associations were linked to proximal breeding locations, we would have expected birds in the same community to have close breeding associations. While this study is limited by a small sample size, the fact that three of the sparrows were from the same social community indicates that social relationships during winter do not carry over to the breeding season.

## Acknowledgments

We would like to thank the many volunteers and interns who helped gather field data and capture (and recapture, especially) the golden-crowned sparrows. Special thanks to Jenny Anderson and Inger Marie Laursen for all their help with field work. We would also like to thank several anonymous reviewers for their comments and feedback.

